# Genetic costs of domestication and improvement

**DOI:** 10.1101/122093

**Authors:** Brook T. Moyers, Peter L. Morrell, John K. McKay

## Abstract

The ‘cost of domestication’ hypothesis posits that the process of domesticating wild species can result in an increase in the number, frequency, and/or proportion of deleterious genetic variants that are fixed or segregating in the genomes of domesticated species. This cost may limit the efficacy of selection and thus reduce genetic gains in breeding programs for these species. Understanding when and how deleterious mutations accumulate can also provide insight into fundamental questions about the interplay of demography and selection. Here we describe the evolutionary processes that may contribute to deleterious variation accrued during domestication and improvement, and review the available evidence for ‘the cost of domestication’ in animal and plant genomes. We identify gaps and explore opportunities in this emerging field, and finally offer suggestions for researchers and breeders interested in understanding or avoiding the consequences of an increased number or frequency of deleterious variants in domesticated species.

## INTRODUCTION

Recently, we have seen a resurgence of evolutionary concepts applied to the domestication and improvement of plants and animals (e.g. Walsh 2007; Wang *et al.* 2014; Gaut, Díez, and Morrell 2015; Kono *et al.* 2016). One particular wave of this resurgence proposes a general ‘cost of domestication’: that the evolutionary processes experienced by lineages during domestication are likely to increase the number of deleterious variants in the genome. This ‘cost’ was first hypothesized by Lu *et al.* (2006), who found an increase in nonsynonymous substitutions, particularly radical amino acid changes, in domesticated compared to wild lineages of rice. These putatively deleterious variants were negatively correlated with recombination rate, which the authors interpreted as evidence that they hitchhiked along with the targets of artificial selection (Figure 1B). Lu *et al.* (2006) conclude that: “The reduction in fitness, or the genetic cost of domestication, is a general phenomenon.” Here we address this claim by examining the evidence that has emerged in the last decade on deleterious variants in domesticated species.

**Figure 1.**
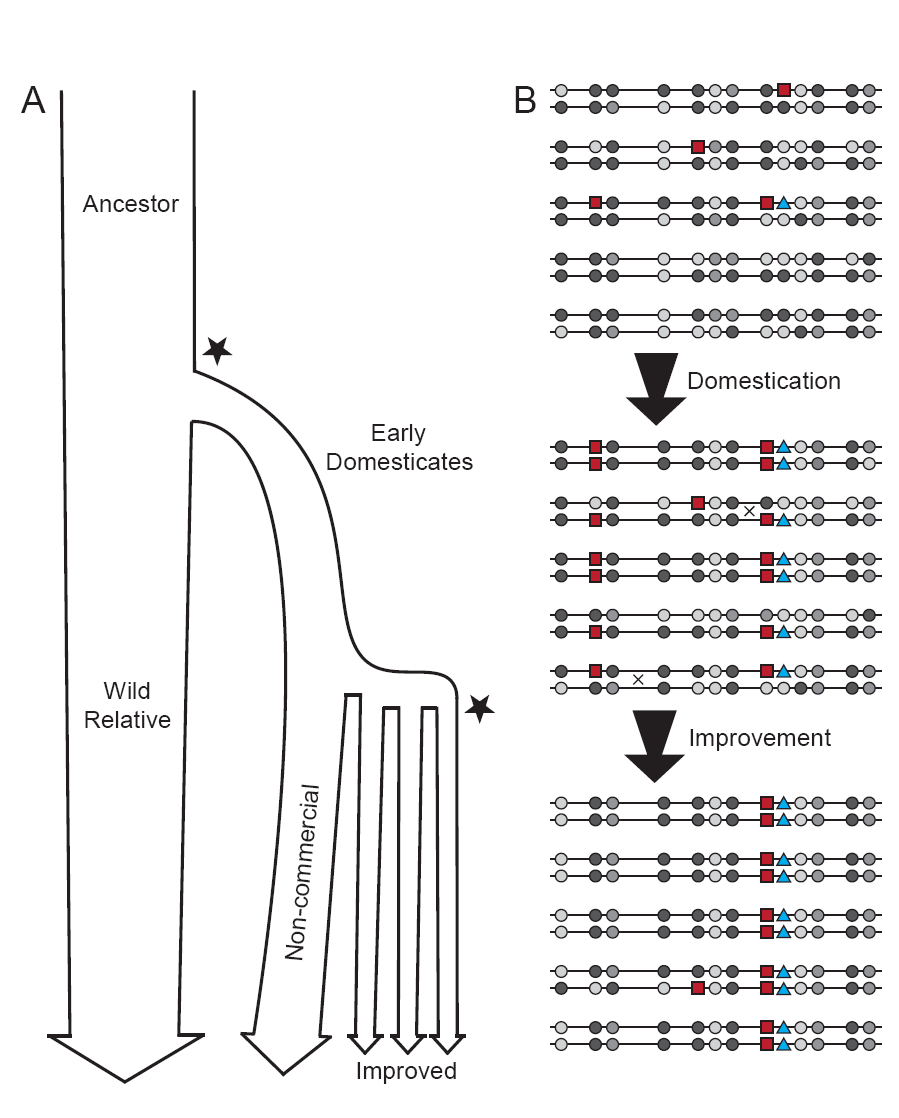
Processes of domestication and improvement. (A) Typical changes in effective population size through domestication and improvement. Stars indicate genetic bottlenecks. These dynamics can be reconstructed by examining patterns of genetic diversity in contemporary wild relative, domesticated noncommercial, and improved populations. (B) Effects of artificial selection (targeting the blue triangle variant) and linkage disequilibrium on deleterious (red squares) and neutral variants (grey circles, shades represent different alleles). In the ancestral wild population, four deleterious alleles are at relatively low frequency (mean = 0.10) and heterozygosity is high (H_O_ = 0.51). After domestication, the selected blue triangle and linked variants increase in frequency (three remaining deleterious alleles, mean frequency = 0.46), heterozygosity decreases (H_O_ = 0.35), and allelic diversity is lost at two sites. Recombination may change haplotypes, especially at sites less closely linked to the selected allele (Xs). After improvement, further selection for the blue triangle allele has: lowered heterozygosity (H_O_ = 0.08), increased deleterious variant frequency (two remaining deleterious alleles, mean = 0.55), and lost allelic diversity at six additional sites.

The processes of domestication and subsequent breeding potentially impose a number of evolutionary effects on populations (Box 1). New mutations can have a range of effects on fitness, from lethal to beneficial. Deleterious variants constitute the mostly directly observable, and likely the most important, source of mutational and therefore genetic load in a population (Box 2). The shape of the distribution of fitness effects of new mutations is difficult to estimate, and estimates suggest it varies across populations and species (Keightley and Eyre-Walker 2007). However, theory predicts that a large proportion of new mutations, particularly those that occur in coding portions of the genome, will be deleterious at least in some proportion of the environments that a species occupies (Ohta 1972; 1992; Gillespie 1994). These predictions are supported by experimental responses to artificial selection and mutation accumulation experiments, as well as molecular genetic studies of variation in populations (Keightley and Lynch 2003; Eyre-Walker, Woolfit, and Phelps 2006; Boyko *et al.* 2008; Kim, Huber, and Lohmueller 2017)

The rate of new mutations in eukaryotes varies, but is likely at least 1 × 10^−8^ / base pair / generation (Baer, Miyamoto, and Denver 2007). For the average eukaryotic genome, individuals should thereby be expected to carry a small number of new mutations not present in the parent genome(s) (Agrawal and Whitlock 2012). The realized distribution of fitness effects for these mutations in a population will be influenced by inbreeding and by effective population size (Gillespie 1999; Whitlock 2000; Arunkumar *et al.* 2015; Keightley and Eyre-Walker 2007). Generally, for a given distribution of fitness effects for segregating variants, we expect to observe relatively fewer strongly deleterious variants and more weakly deleterious variants in smaller populations and in populations with higher rates of inbreeding (Figure 2A).

**Figure 2.**
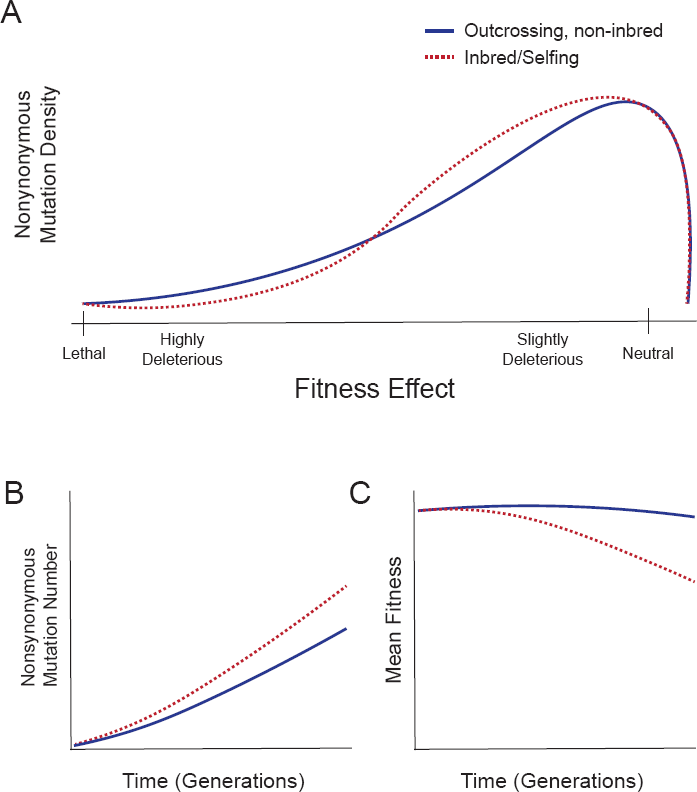
Effects of inbreeding on mutations and fitness. (A) Theoretical density plot of fitness effects for segregating variants. In outcrossing, non-inbred populations (solid navy line), more variants with highly deleterious, recessive effects persist that are rapidly exposed to selection and purged in inbred populations (dotted red line), shifting the left side of the distribution towards more neutral effects. At the same time, the reduced effective population size created through inbreeding causes on average higher loss of slightly advantageous mutations and retention of slightly deleterious mutations, shifting the right side of the distribution towards more deleterious effects. (B) Accumulation of nonsynonymous mutations is accelerated in selfing individuals (dotted red line) relative to outcrossing individuals (solid navy line). Synonymous mutations accumulate at similar rates in both populations. Adapted from simulation results in Arunkumar *et al.* 2015. (C) As a consequence of (B), mean individual fitness drops in selfing relative to outcrossing populations. Adapted from Arunkumar *et al.* 2015.

Domestication and improvement often involve increased inbreeding. Allard (1999) pointed out that many important early cultigens were produced by inbreeding, with inbred lines offering agronomic and morphological consistency in the crop. In some cases, domestication involved a switch in mating system from outcrossing to highly selfing (e.g. in rice; Kovach, Sweeney, and McCouch 2007). The practice of producing inbred lines, often through single-seed descent, with the goal of reducing heterozygosity and creating genetically ‘stable’ varieties, remains a major activity in plant breeding programs. Reduced effective population sizes (N_e_) during domestication and artificial selection on favorable traits also constitute forms of inbreeding (Figure 1). Even in species without the capacity to self-fertilize, inbreeding that results from selective breeding can dramatically change patterns of genomic variation. For example, the first sequenced dog genome, from a female boxer, has runs of homozygosity that span 62% of genome, with an N50 length of 6.9 Mb (versus heterozygous region N50 length 1.1 Mb; Lindblad-Tor *et al.* 2005). This level of homozygosity drastically reduces the effective recombination rate, as crossover events often exchange identical chromosomal segments. The likelihood of a beneficial allele moving into a genomic background with fewer linked deleterious alleles is thus reduced.

Linked selection, or interference among mutations, has much the same effect. For a trait influenced by multiple loci, interference between linked loci can limit the response to selection (Hill and Robertson 1966). Deleterious variants are more numerous than those that have a positive effect on a trait and thus should constitute a major limitation in responding to selection (Felsenstein 1974). This forces selection to act on the net effect of favorable and unfavorable mutations in linkage (Figure 1B). As homozygosity and LD increase, N_e_ decreases, and so selection is less effective at purging moderately deleterious mutations and new slightly beneficial mutations are more likely to be lost to genetic drift (sometimes termed ‘inbreeding depression’). These dynamics will shift the distribution of fitness effects for segregating variants and result in greater accumulation of deleterious variation over time (Figure 2A-C).

Overall, the cost of domestication hypothesis posits that compared to their wild relatives domesticated lineages will have:

1. Deleterious variants at higher number, frequency, and/or proportion
2. Enrichment of deleterious variants in linkage disequilibrium with loci subject to strong, positive, artificial selection

Among domesticated species, these effects may differ between lineages that experienced domestication only (e.g. landraces and non-commercial populations) and those that were subject to modern improvement (e.g. ‘elite’ varieties and commercial breeds). Domestication typically involves a genetic bottleneck followed by a long period of relatively weak and possibly varying selection, while during the process of improvement, intense selection over short time periods is coupled with limited recombination and an additional reduction in N_e_, often followed by rapid population expansion (Figure 1A; Yamasaki, Wright, and McMullen 2007). Gaut, Díez, and Morrell (2015) propose that elite crop lines could harbor a lower proportion of deleterious variants relative to landraces due to strong selection for yield during improvement, but the opposing pattern could be driven by lower N_e_ and thus increased genetic drift, limited effective recombination, and rapid population expansion. At least one study, in sunflower, shows little difference in the composition of deleterious variants between landrace and elite lines (Renaut and Rieseberg 2015). It is likely that the relative influence of these factors varies dramatically across domesticated systems.

### Box 1. Domesticated lineages may experience

- Increased number of deleterious variants:

⃘ Deleterious mutations may accumulate at higher rates in domesticated lineages versus their wild relatives due to the reduced efficacy of selection relative to genetic drift. Mutations that would be purged with a larger N_e_ or with higher effective recombination rates are instead retained. This is reflected in a shift in the distribution of fitness effects for segregating variants towards more moderately deleterious alleles (Figure 2A).
⃘ **However:** any significant increase in inbreeding, particularly the transition from outcrossing to selfing, can result in the purging of recessive, highly deleterious alleles, as these alleles are exposed in homozygous genotypes more frequently (Arunkumar *et al.* 2015). This shifts the more deleterious end of the distribution of fitness effects for segregating variants towards neutrality (Figure 2A).

- Increased frequency of deleterious variants:

⃘ As above, reduced N_e_ (due to inbreeding, genetic bottlenecks, or strong selection) or reduced effective recombination rate will increase the strength of genetic drift relative to selection. Stronger genetic drift can allow deleterious variants to reach higher frequencies. In the case of inbreeding, this pattern is called inbreeding depression, but it can occur whenever N_e_ decreases.
⃘ Deleterious variants that are in linkage disequilibrium with a target of artificial selection can also increase in frequency through genetic hitchhiking, as long as their fitness effects are smaller than the strength of selection on the targeted variant (Figure 1B; Hartfield and Otto 2011; Assaf, Petrov, and Blundell 2015).
⃘ The rapid population expansion, coupled with long-distance migration common to the demographic history of many domesticated lineages, can result in the accumulation of deleterious genetic variation known as ‘expansion load’ (Peischl *et al.* 2013; Lohmueller 2014). This occurs when serial bottlenecks are followed by large population expansions (e.g. large local carrying capacities, large selection coefficients, long distance dispersal; Peischl *et al.* 2013). This phenomenon is due to the accumulation of new deleterious mutations at the ‘wave front’ of expanding populations, which then rise to high frequency via drift regardless of their fitness effects (also known as ‘gene surfing’, or ‘allelic surfing’ Klopfstein, Currat, and Excoffier 2006; Travis *et al.* 2007).

### Box 2: What is the link between deleterious variants and the long-standing concept of genetic load?

The classic concept of genetic load identifies a reduction in fitness from a mean or optimal genotype (Haldane 1937). Deleterious variants are probably the largest single contributor to genetic load via ‘mutational load’ (Muller 1950; Felsenstein 1974; Agrawal and Whitlock 2012). Due to this, the term ‘genetic load’ has often been used colloquially to indicate mutational load, and attempts to estimate genetic load have involved identifying and quantifying deleterious genetic variants. However, the genetic load of an individual is not merely a count of deleterious variants, but is dependent on other factors, including the distributions of fitness effects and dominance coefficients for deleterious variants (see Henn *et al.* 2015). These factors are difficult to quantify and estimates of genetic load are sensitive to their values (Boyko *et al.* 2008; Lohmueller *et al.* 2008; Lohmueller *et al.* 2014; Simons *et al.* 2014). For this reason, much of the modern literature has eschewed discussion of genetic load and focused on the various means of counting the number of deleterious variants (see Chun and Fay 2009; Marth *et al.* 2011). Similarly, this review primarily addresses patterns of deleterious variation in domesticated species, with the view that deleterious variants likely contribute significantly to genetic load in many cases.

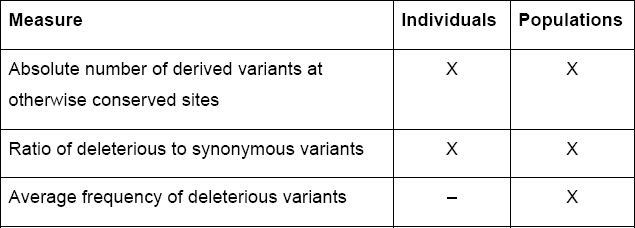
Measures used to quantify the difference in mutational load among individuals and populations include:

| Measure | Individuals | Populations |
| --- | --- | --- |
| Absolute number of derived variants at otherwise conserved sites | X | X |
| Ratio of deleterious to synonymous variants | X | X |
| Average frequency of deleterious variants | – | X |

### Box 3: Definitions

*allelic (or gene) surfing:* the increase in frequency of a genetic variant at the wave front of an expanding population

*ancient DNA:* DNA samples from archaeological or historical remains

*deleterious variant:* a genetic variant that reduces fitness

*derived mutation:* a genetic variant that involves a change from the ancestral state as inferred from a related, outgroup taxon or from sampling of the ancestral genome (e.g. using ancient DNA)

*effective population size* (N_e_): the number of individuals contributing offspring to the next generation of a population, versus the census population size

*genetic bottleneck:* a large reduction in census population size

*genetic hitchhiking:* when a genetic variant increases in frequency owing to linkage disequilibrium with a variant subject to positive selection

*haplotype:* set of genetic variants in LD with each other

*linkage disequilibrium (LD):* nonrandom association of alleles at two or more loci

*LD N50:* the approximate physical or genetic distance over which LD decays to half of its maximum value

*linked selection:* the tendency for selection on a variant to affect the frequency of nearby variants due to LD

*private alleles:* genetic variants unique to a population or set of populations

*runs of homozygosity:* regions of the genome where both of a pair of chromosomes are highly similar, thus increasing LD due to reduced effective recombination

*site frequency spectrum:* the distribution of variant frequencies in a population

## Deleterious variation in Domesticated Species

The rapidly expanding number of genome-wide datasets has made the study of deleterious variants increasingly feasible. Since Lu *et al.* (2006) first hypothesized the accumulation of deleterious variants as a ‘cost of domestication’ in rice, similar studies have examined evidence for such costs in other crops and domesticated animals.

A number of approaches have been used to quantify the composition and effect of deleterious variants in domesticated genomes. These approaches largely parallel those used in studies of the effects of demography on human populations (Lohmueller 2014; Henn *et al.* 2015). Some studies have inferred a cost of domestication indirectly from the accumulation of a larger proportion or number of nonsynonymous variants in domesticated lineages versus their wild relatives, assuming that nonsynonymous changes are on average deleterious (e.g. Lu *et al.* 2006; Cruz, Vilà, and Webster 2008). Other studies have looked for reduced genetic diversity or longer distance linkage disequilibrium (e.g. Lindblad-Toh *et al.* 2005; Lam *et al.* 2010), and therefore reduced effective recombination and presumably reduced efficacy of selection in the same comparison. These approaches assume that wild relatives of domesticated species have not themselves experienced population bottlenecks, shifts in mating system, or any of the other processes that could affect patterns of deleterious variation relative to their shared ancestors.

A more direct approach used to estimate the cost of domestication is to assess the number and proportion of putatively deleterious variants present in populations of domesticated species. The measures used include 1) the absolute number of variants at derived sites, 2) the ratio of deleterious to synonymous variants, and 3) an increase in the frequency of deleterious variants within a population (Table S1; Box 2; also reviewed in Lohmueller 2014). We discuss approaches to identifying deleterious variants in more depth below.

Several approaches attempt to translate the number of observed deleterious variants into an estimate of genetic load. The first approach is to assume that the degree of phylogenetic constraint on a variant provides an estimate of the strength of purifying selection. Summing over the constraint scores (typically using the GERP++ software; Cooper et al. 2005) relative to the number of deleterious variants provides a means of estimating mutational load (e.g. Wang *et al. bioRxiv;* Mardsen *et al.* 2016). A more general translation of deleterious variants into genetic load depends on three factors: the distribution of fitness effects of deleterious variants, a model of either additive or multiplicative effects on fitness, and an estimate of dominance of deleterious variants (Arunkumar *et al.* 2015; Henn *et al.* 2015; Henn *et al.* 2016; Brandvain and Wright 2016). This approach directly addresses the question of genetic load, but does not tell us whether the load is the result of domestication or other evolutionary processes (unless wild relatives are also assayed, again assuming that their own evolutionary trajectory has not simultaneously been affected). It also requires accurate, unbiased algorithms for identifying deleterious variants from neutral variants (see below).

Studies taking these approaches in domesticated species are presented qualitatively in Table 1 (and quantitatively in Supplementary Table 1). We searched the literature using Google Scholar with the terms (“genetic load” or “deleterious”) and “domesticat*”, and in many cases followed references from one study to the next. To the best of our knowledge, the studies in Table 1 represent the majority of the extant literature on this topic. We excluded studies examining only the mitochondrial or other non-recombining portions of the genome. In a few cases where numeric values were not reported, we extracted values from published figures using relative distances as measured in the image analysis program ImageJ (Schneider, Rasband, and Eliceiri 2012), or otherwise extrapolated values from the provided data. Where exact values were not available, we provide estimates. Please note that no one study (or set of genotypes) contributed values to all columns for a particular domestication event, and that in many cases methods used or statistics examined varied across species.

**Table 1.**
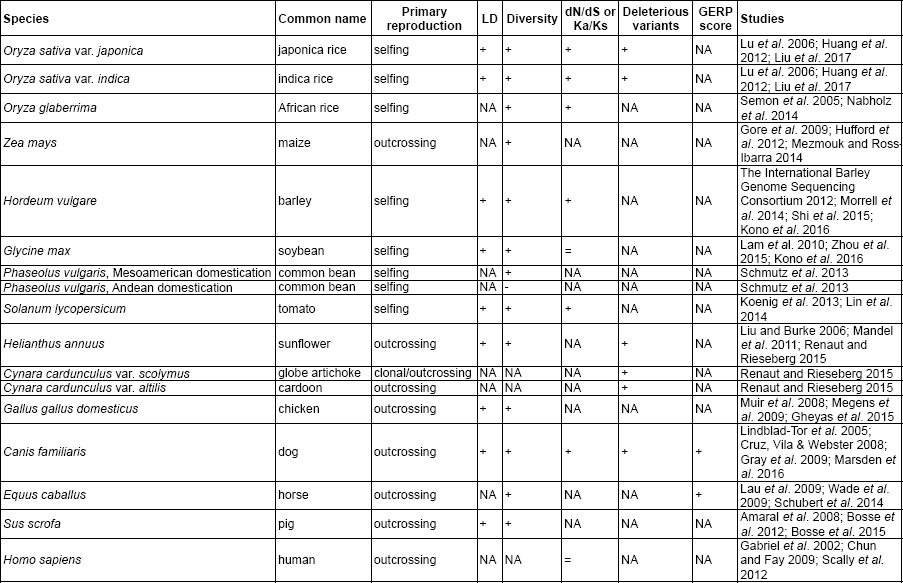
Table 1. Evidence for a cost of domestication across domesticated plants and animals, with *Homo sapiens* included for comparison. The first three columns identify the taxa and major form of propagation and reproduction. Following columns indicate whether data from the domesticated lineage versus wild relatives or ancestors fits (+), is equivocal (=) or goes against (-) expectations for the ‘cost of domestication’ hypothesis, with those expectations being: extended linkage disequilibrium (LD), reduced genetic diversity (Diversity), higher rate or number of nonsynonymous to synonymous substitutions or variants (dN/dS, Ka/Ks, counts), more or higher frequency deleterious variants (Deleterious variants), or higher estimated genetic load (GERP score). The values for each of these and additional data are available in Supplementary Table 1.

### Recombination and linkage disequilibrium

The process of domestication may increase recombination rate, as measured in the number of chiasmata per bivalent. Theory predicts that recombination rate should increase during periods of rapid evolutionary change (Otto and Barton 1997), and in domesticated species this may be driven by strong artificial selection or small *N_e_s* (Otto and Barton 2001). This is supported by the observation that recombination rates are higher in many domesticated species compared to wild relatives (e.g. chicken, Groenen *et al.* 2009; honey bee, Wilfert *et al.* 2007; and a number of cultivated plant species, Ross-Ibarra 2004; but not mammals, Muñoz-Fuentes *et al.* 2014). In contrast, recombination in some domesticated species is limited by genomic structure (e.g. in barley, where 50% of physical length and many functional genes are in (peri-)centromeric regions with extremely low recombination rates; The International Barley Genome Sequencing Consortium 2012), although this structure may be shared with wild relatives. Even when actual recombination rate (chiasmata per bivalent) increases, effective recombination may be reduced in many domesticated species due to an increase in linkage disequilibrium (LD) due to runs of homozygousity. In other words, chromosomes may physically recombine, but if the homologous chromosomes contain identical sequences then no ‘effective recombination’ occurs and the outcome is no different than no recombination. In a striking example, the effective recombination rate in maize populations has decreased by an estimated 83% compared to that in the wild relative teosinte (Wright *et al.* 2005; with comparable estimates in Hufford *et al.* 2012). We note that Mezmouk and Ross-Ibarra (2014) found that deleterious variants are not enriched in areas of low recombination in maize (although see McMullen *et al.* 2009; Rodgers-Melnick *et al.* 2015).

Strong directional selection like that imposed during domestication can reduce genetic diversity in chromosomal regions linked to the selected locus, creating runs of homozygousity (Sved 1971; Maynard Smith and Haigh 1974). These regions of extended LD are among the signals used to identify targets of selection and reconstruct the evolutionary history of domesticated species (e.g. Tian, Stevens, and Buckler 2009). Any increase in inbreeding (e.g. after a population bottleneck or a shift in mating system from outcrossing to selfing) will have the same effect across the entire recombining portion of the genome, as heterozygosity decreases with each generation (Pritchard and Przeworski 2001; Charlesworth 2003). Extended LD has consequences for the efficacy of selection: deleterious variants linked to larger-effect beneficial alleles can no longer recombine away, and beneficial variants that are not in LD with larger-effect beneficial alleles may be lost by genetic drift (Hartfield and Otto 2011; Assaf, Petrov, and Blundell 2015). In our review of the literature, LD decays most rapidly in outcrossing plant species (maize and sunflower), and extends much further in self-fertilizing plants and in domesticated animals (Table S1). In all cases where we have data for wild relatives, LD decays more rapidly in wild lineages than in domesticated lineages (Table S1).

While we primarily report mean LD in Table S1, linkage disequilibrium also varies among varieties and breeds of the same species. For example, in domesticated pig breeds the length of the genome covered by runs of homozygosity, which extend LD, ranges from 13.4 to 173.3 Mb, or 0.5-6.5% (Traspov *et al.* 2016; Box 3). Similarly, among dog breeds average LD decay (to r^2^ ≤ 0.2) ranges from 20 kb to 4.2 Mb (Gray *et al.* 2009). In both species, the decay of LD for wild individuals occurs over shorter distances (Table S1). This suggests that patterns of LD in these species have been strongly impacted by breed-specific demographic history (i.e. the process of improvement), in addition to the shared process of domestication.

### Genetic diversity

We see consistent loss of genetic diversity when ‘improved’ or ‘breed’ genomes are compared to domesticated ‘landrace’ or ‘non-commercial’ genomes, and again when domesticated genomes are compared to the genomes of wild relatives (Table 1; Table S1). This ranges from ~5% nucleotide diversity lost between wolf populations and domesticated dogs (Gray *et al.* 2009) to 77% lost between wild and improved tomato populations (Lin *et al.* 2014). The only case where we see a gain in genetic diversity is in the Andean domestication of the common bean, where gene flow with the more genetically diverse Mesoamerican common bean is likely an explanatory factor (Schmutz *et al.* 2013). This pattern is consistent with a broader review of genetic diversity in crop species: Miller and Gross (2011) found that annual crops had lost an average of ~40% of the diversity found in their wild relatives. This same study found that perennial fruit trees had lost an average of ~5% genetic diversity, suggesting that the impact of domestication on genetic diversity is strongly influenced by life history (see also Gaut, Díez, and Morrell 2015). Given a relatively steady evolutionary trajectory for wild populations, loss of genetic diversity in domesticated populations can be attributed to artificial selection or reduced *N_e_* due to increased inbreeding and genetic bottlenecks. On an evolutionary timescales, even the oldest domestications occurred recently relative to the rate at which new mutations can recover the loss. This is especially true for non-recombining portions of the genome like the mitochondrial genome or sex chromosomes. Most modern animal breeding programs are strongly sex-biased, with few male individuals contributing to each generation. In horses, for example, this has likely led to almost complete loss of polymorphism on the Y chromosome via genetic drift (Lippold *et al.* 2011). Loss of allelic diversity can reduce the efficacy of selection by reducing additive genetic variance within species (Eyre-Walker, Woolfit, and Phelps 2006). However, a loss of genetic diversity alone does not necessarily signal a corresponding increase in the frequency or proportion of deleterious variants, and so is not sufficient evidence of a cost of domestication.

### Synonymous versus nonsynonymous variation

In six species, the domesticated lineage shows an increase in genome-wide nonsynonymous to synonymous substitution rate or number compared to a wild lineage (Table 1). This is also true across the domesticated lineages of cattle (especially the domesticated cow; MacEachern *et al.* 2009). An exception is in soybean, where the domesticated *Glycine max* and wild *G. soja* genomes contain approximately the same proportion of nonsynonymous to synonymous single nucleotide polymorphisms (Lam *et al.* 2010), which are not directly comparable to substitutions but should exhibit similar patterns under the cost of domestication hypothesis. These differences in nonsynonymous substitution rate are likely driven by differences in N_e_ (Eyre-Walker and Keightley 2007; Woolfit 2009). If nonsynonymous mutations are on average deleterious, as theory and empirical data suggest (Keightley and Lynch 2003; Eyre-Walker, Woolfit, and Phelps 2006; Boyko *et al.* 2008; Kim, Huber, and Lohmueller 2017), then an increase in nonsynonymous substitutions will reduce mean fitness (Figure 2B-C). The comparisons in Table 1 suggest that this has occurred in domesticated species. This result differs from Moray, Lanfear, and Bromham (2014), who examined rates of mitochondrial genome sequence evolution in domesticated animals and their wild relatives and found no such consistent pattern. This difference may be attributable to the focus of each review (genome-wide versus mitochondria) or, as the authors speculate, to genetic bottlenecks in some of the wild relatives included in their study.

The ratio of nonsynonymous to synonymous substitutions may not be a good estimate for mutational load. For one, nonsynonymous substitutions are particularly likely to have phenotypic effects (Kono *et al. bioRxiv*). Variants annotated as deleterious based on sequence conservation (see below) can in some cases contribute to agronomically important phenotypes (e.g. Nie *et al.* 2015; see also Albalat and Cañestro 2016), and artificial selection during domestication and improvement is likely to drive a portion of these variants associated with favorable phenotypes to higher frequencies (or fixation) in domesticated lineages (Kono *et al.* 2016). In addition, estimates of the proportion of nonsynonymous sites with deleterious effects range from 0.03 (in bacteria; Hughes 2005) to 0.80 (in humans; Fay *et al.* 2001; The Chimpanzee Sequencing and Analysis Consortium 2005). It follows that nonsynonymous substitution rate may have a poor correlation with patterns of deleterious variation and mutational load, at least at larger taxonomic scales. However, the consensus proportion of deleterious variants in Table S1 is between 0.05 and 0.25, which spans a smaller range. Finally, dN/dS and related ratios may be flawed estimates of functional divergence because they rely on the assumption that synonymous mutations are neutral and thus can control for substitution rate variation. This assumption may not hold true, especially for closely related taxa (e.g. Wolf *et al.* 2009; see also Kryazhimskiy and Plotkin 2008), and at least one study shows elevated rates of synonymous relative to non-coding substitutions in domesticated lineages (MacEachern *et al.* 2009).

### Deleterious variants

An increase in mutational load can come from an increased frequency or number of deleterious variants. The first can be assessed by examining shared deleterious variants between wild and domesticated lineages, and both by comparing shared and private deleterious variants across lineages. Looking at deleterious variants that are at high frequency across all domesticated varieties may provide insight into the early processes of domestication, while looking at deleterious variants with varying frequencies among domesticated varieties may provide insight into the processes of improvement. We recommend that researchers studying deleterious variants report results per genome, and then compare across genomes and among lineages. Many, but not all, of the studies in Table 1 take this approach. The total number or frequency of deleterious variants within a population will necessarily depend on the size of that population, and so sufficient sampling is important before values can be compared across populations. Values per genome are easier to compare across most available genomic datasets.

Renaut and Rieseberg (2015) found a significant increase in both shared and private deleterious mutations in domesticated relative to wild lines of sunflower, and similar patterns in two additional closely-related species: cardoon and globe artichoke (Table 1). This pattern also holds true for *japonica* and *indica* domesticated rice (Liu *et al.* 2017) and for the domesticated dog (Marsden *et al.* 2016) compared to their wild relatives (Table 1). In horses, deleterious mutation load as estimated using Genomic Evolutionary Rate Profiling (GERP) appears to be higher in both domesticated genomes and the extant wild relative (Przewalski's horse, which went through a severe genetic bottleneck in the last century) compared to an ancient wild horse genome (Schubert *et al.* 2014). These four studies, spanning a wide taxonomic range, suggest that an increase in the number and proportion of deleterious variants may be a general consequence of domestication. However, the other studies we present that examined deleterious variants in domesticated species did not include any sampling of wild lineages. Without sufficient sampling of a parallel lineage that did not undergo the process of domestication, it is difficult to assess whether the ‘cost of domestication’ is indeed general.

### Identifying dangerous hitchhikers

The effect size of a deleterious mutation is negatively correlated with its likelihood of increasing in frequency through any of the mechanisms we discuss here. That is, variants with a strongly deleterious effect are more likely to be purged by selection than mildly deleterious variants. Similarly, mutations that have a consistent, environmentally-independent deleterious effect are more likely to be purged than mutations with environmentally-plastic effects. In the extreme case, we would never expect mutations that have a consistently lethal effect in a heterozygous state to contribute to a persistent cost of domestication, as these would be lost in the first generation of their appearance in a population. However, mutations with consistent, highly deleterious effects are likely rare relative to those with smaller or environment-dependent effects, especially in inbred populations (Figure 2A; Arunkumar *et al.* 2015). When thinking about patterns of deleterious variation, it is therefore important to recognize that the effect of any particular mutation can depend on context, including genomic background and developmental environment. This complexity likely makes these classes of deleterious variants more difficult to identify, and we might not expect these variants to show up in bioinformatic screens (discussed below). Furthermore, all of these expectations are modified by linkage: when deleterious variants are in LD with targets of artificial selection, they are more likely to evade purging even with consistent, large, deleterious effects (Figure 1B).

We searched the literature for examples of specific deleterious variants that hitchhiked along with targets of selection during domestication and improvement. We did not find many such cases, so we describe each in detail here. The best-characterized example comes from rice, where an allele that negatively affects yield under drought *(qDTY1.1)* is tightly linked to the major green revolution dwarfing allele *sd1* (Vikram *et al.* 2015). Vikram *et al.* (2015) found that the *qDTY1.1* allele explained up to 31% of the variance in yield under drought across three RIL populations and two growing seasons. Almost all modern elite rice varieties carry the *sd1* allele (which increases plant investment in grain yield), and as a consequence these varieties are highly sensitive to drought. The discovery of the *qDTY1.1* allele has enabled rice breeders to finally break the linkage and create drought tolerant, dwarfed lines.

In sunflower, the *B* locus affects branching and was a likely target of selection during domestication (Bachlava *et al.* 2010). This locus has pleiotropic effects on plant and seed morphology that, in branched male restorer lines, mask the effect of linked loci with both ‘positive’ (increased seed weight) and ‘negative’ (reduced seed oil content) effects (Bachlava *et al.* 2010). To properly understand these effects required a complex experimental design, where these linked loci were segregated in unbranched (*b*) and branched (*B*) backgrounds. Managing these effects in the heterotic sunflower breeding groups has likely also been challenging.

A similarly complex narrative has emerged in maize. The gene *TGA1* was key to evolution of ‘naked kernels’ in domesticated maize from the encased kernels of teosinte (Wang *et al.* 2015). This locus has pleiotropic effects on kernel features and plant architecture, and is in linkage disequilibrium with the gene *SU1,* which encodes a starch debranching enzyme (Brandenburg *et al.* 2017). *SU1* was targeted by artificial selection during domestication (Whitt *et al.* 2002), but also appears to be under divergent selection between Northern Flints and Corn Belt Dents, two maize populations (Brandenburg *et al.* 2017). This is likely because breeders of these groups are targeting different starch qualities, and this work may have been made more difficult by the genetic linkage of *SU1* with *TGA1.*

In the above cases, the linked allele(s) with negative agronomic effects are unlikely to be picked up in a genome-wide screen for deleterious variation, as they are segregating in wild or landrace populations and are not necessarily disadvantageous in other contexts. One putative ‘truly’ deleterious case is in domesticated chickens, where a missense mutation in the thyroid stimulating hormone receptor *(TSHR)* locus sits within a shared selective sweep haplotype (Rubin *et al.* 2010). However, Rubin *et al.* (2010) argue this is more likely a case where a ‘deleterious’ (i.e. non-conserved) allele was actually the target of artificial selection and potentially contributed to the trait of year-round egg laying in chickens.

In the Roundup Ready (event 40-3-2) soybean varieties released in 1996, tight linkage between the transgene insertion event and another allele (or possibly an allele created by the insertion event itself) reduced yield by 5-10% (Elmore *et al.* 2001). This is not quite genetic hitchhiking in the traditional sense, but the yield drag effect persisted through backcrossing of the transgene into hundreds of varieties (Benbrook 1999). This effect likely explains, at least in part, why transgenic soybean has failed to increase realized yields (Xu *et al.* 2013). Consistent with this prediction, a second, independent insertion event in Roundup Ready 2 Yield® does not suffer from the same yield drag effect (Horak *et al.* 2015).

It is likely that ‘dangerous hitchhiker’ examples exist that have either gone undetected by previous studies (possibly due to low genomic resolution, limited phenotyping, or limited screening environments), have been detected but not publicized, or are buried among other results in, for example, large QTL studies. It is also possible that the role of genetic hitchhiking has not been as important in shaping genome-wide patterns of deleterious variation as previously assumed.

## Gaps and Opportunities

*How much of the genome is in LD with major targets of artificial selection?* Artificial selection during domestication targets a clear change in the optimal multivariate phenotype. This likely affects a significant portions of the genome: available estimates include 2–4% of genes in maize (Wright *et al.* 2005) and 16% of the genome in common bean (Papa *et al.* 2007) targeted by selection during domestication. For crops, traits such as seed dormancy, branching, indeterminate flowering, stress tolerance, and shattering are known to be selected for different optima between artificial versus natural selection (Takeda and Matsuoka 2008; Gross and Olsen 2010). In some cases, we know the loci that underlie these domestication traits. One well-studied example is the green revolution dwarfing gene, *sd1* in rice. *sd1* is surrounded by a 500kb region (~13 genes) with reduced allelic diversity in *japonica* rice (Asano *et al.* 2011). Another example in rice is the *waxy* locus, where a 250 kb region shows reduced diversity consistent with a selective sweep in temperate *japonica* glutinous varieties (Olsen *et al.* 2006). The difference in the size of the region affected by these two selective sweeps may be because the strength of selection on these two traits varied, with weaker selection at *waxy* then *sd1.* Unfortunately, the relative strength of selection on domestication traits is largely unknown, and other factors can also influence the size of the genomic region affected by artificial selection. The physical position of selected mutation can have large effect on this via gene density and local recombination rate (e.g. in rice, Flowers *et al.* 2012). This explanation has been invoked in maize (Wright *et al.* 2005), where the extent of LD surrounding domestication loci is highly variable. For example a 1.1 Mb region (~15 genes) lost diversity during a selective sweep on chromosome 10 in maize (Tian, Stevens, and Buckler 2009), but only a 60–90 kb extended haplotype came with the *tb1* domestication allele (Clark *et al.* 2004). While these case studies provide examples of sweeps resulting from domestication, they also show that the size of the affected region is highly variable, and we don’t yet know how this might impact patterns of deleterious variation. This is compounded by the fact that even in highly researched species we don’t always know the number or location of genomic targets of selection during domestication (e.g. in maize; Hufford *et al.* 2012), especially if any extended LD driven by artificial selection has eroded or the intensity or mode of artificial selection has changed over time.

*What’s “worse” in domesticated species: hitchhikers, drifters, or inbreds?* We do not have a clear sense of which evolutionary processes contribute most to the putative cost of domestication, and sometimes see contrasting patterns across species. In maize, relatively few putatively deleterious alleles are shared across all domesticated lines (hundreds vs. thousands; Mezmouk and Ross-Ibarra 2014), which points to a larger role for the process of improvement than for domestication in driving patterns of deleterious variation. In contrast, the increase in dog dN/dS relative to wolf populations appears to not be driven by recent inbreeding (i.e. improvement) but by the ancient domestication bottleneck common to all dogs (Marsden *et al.* 2016).

Of the deleterious alleles segregating in more than 80% of maize lines, only 9.4% show any signal of positive selection (Mezmouk and Ross-Ibarra 2014). This suggests that hitchhiking during domestication played a relatively small role in the increased frequency of deleterious variants in maize. The same study found little support for enrichment of deleterious SNPs in areas of reduced recombination (Mezmouk and Ross-Ibarra 2014). However, Rodgers-Melnick *et al.* (2015) present contrasting evidence supporting enrichment of deleterious variants in regions of low recombination, and the authors argue that this difference is due to the use of a tool that does not rely on genome annotation (Genomic Evolutionary Rate Profiling, or GERP; Cooper *et al.* 2005). This complex narrative has emerged from just one well-studied domesticated species, and it is likely that each species will present new and different complexities.

### Specific differences

Examining general differences between wild and domesticated lineages ignores species-specific demographic histories and changes in life history, which may be important contributors to patterns of deleterious variation. Although we found some general patterns (e.g. loss of genetic diversity, Table 1), we also see clear exceptions (e.g. the Andean common bean). We can attribute these exceptions to particular demographic scenarios (e.g. gene flow with Mesoamerican common bean populations), assuming we have sufficient archeological, historical, or genetic data. One clear problem is our inability to sample ancestral (pre-domestication) lineages, and our subsequent reliance on sampling of current wild relative lineages that have their own unique evolutionary trajectories. Sequencing ancient DNA can provide some insight into the history of these lineages and their ancestral states (e.g. in horses; Schubert *et al.* 2014). Currently, we know very little about most domesticated species’ histories. We are still working towards understanding dynamics since domestication: even in highly-researched species like rice, the number of domestication events and subsequent demographic dynamics are hotly contested (Kovach, Sweeney, and McCouch 2007; Gao and Innan 2008; He *et al.* 2011; Molina *et al.* 2011; Huang *et al.* 2012; Gross and Zhao 2014; Civáň *et al.* 2015; Chen, Huang, and Han 2016; Choi *et al.* 2017).

A second challenge, briefly mentioned above, is understanding the relative importance of any one factor in any particular domestication event. Freedman, Lohmueller, and Wayne (2016) provide an in-depth review on this question in the domesticated dog, but it is unclear how general the relative contributions of selection and demography in dogs may be to other species. For example, in rice the shift to selfing from outcrossing during domestication appears to have played a larger role than the domestication bottleneck in shaping deleterious variation (Liu *et al.* 2017). This is useful in understanding rice domestication and its impact, but similar studies would need to be conducted across domesticated species to understand the generality of this dynamic. It is possible that general patterns may be drawn from subsets of domesticated species (e.g. vertebrates versus vascular plants, short-lived versus long-lived, outcrossing versus selfing versus clonally propagated etc.). For one, there may be general differences between annual and perennial crops, including less severe domestication bottlenecks and higher levels of gene flow from wild populations in perennials (Miller and Gross 2011; Gaut, Díez, and Morrell 2015).

### Predictive algorithms

The identification of individual deleterious variants typically relies on sequence conservation. If a variant occurs at a particular nucleotide site or encoded amino acid is invariant across a phylogenetic comparison, then it is putatively deleterious. More advanced approaches use estimates of synonymous substitution rates at a locus to improve estimates of constraint on a nucleotide site (Chun and Fay 2009). The majority of ‘SNP annotation’ approaches are intended for the annotation of amino acid changing variants, although at least two approaches (GERP++: Davydov *et al.* 2010; PHAST: Hubisz, Pollard, and Siepel 2010) can be applied to noncoding sequences when nucleotide sequences can be aligned across species. This estimation of phylogenetic constraint is heavily dependent on the sequence alignment; new annotation approaches have sought to use more consistent sets of alignments across loci. Both GERP++ and MAPP permit users to provide alignments for SNP annotation (Davydov *et al.* 2010; Stone and Sidow 2005). The recently reported tool BAD_Mutations (Kono *et al.* 2016; Kono *et al.* bioRxiv) permits the use of a consistent set of alignments for the annotation of deleterious variants by automating the download and alignment of the coding portion of plant genomes from Phytozome and Ensembl Plants. This currently includes 50+ sequenced angiosperm genomes (Goodstein *et al.* 2010; Kersey *et al.* 2016). Currently, the tool is configured for use with angiosperms, but could be applied to other organisms.

Using phylogenetic conservation may be problematic in domesticated species, as relaxed, balancing, or diversifying selection in agricultural environments could lift constraint on sites under purifying or stabilizing selection in wild environments (assuming some commonalities within these environment types). In one such case, two AGPase small subunit paralogs in maize appear to be under diversifying and balancing selection, respectively, even though these subunits are likely under selective constraint across flowering plants (Georgelis, Shaw, and Hannah 2009; Corbi *et al.* 2010). Similarly, broadly ‘deleterious’ traits may have been under positive artificial selection in domesticated species. The loci underlying these traits could be flagged as deleterious in bioinformatic screens despite increasing fitness in the context of domestication. For example, the *fgf4* retrogene insertion that causes chrondrodysplasia (short-leggedness) in dogs would likely have a strongly deleterious effect in wolves, but has been positively selected in some breeds of dog (Parker *et al.* 2009). Finally, bioinformatic approaches that rely on phylogenetic conservation are likely to miss variants with effects that are only deleterious in specific environmental or genomic contexts (plastic or epistatic effects), or which reduce fitness specifically in agronomic or breeding contexts. Specific knowledge of the phenotypic effects of putatively deleterious mutations is necessary to address these issues, but as we discuss below these data are challenging to obtain.

Information from the site frequency spectrum, or the number of times individual variants are observed in a sample, can provide additional information about which variants are most likely to be deleterious. In resequencing data from many species, nonsynonymous variants typically occur at lower average frequencies than synonymous variants (see Nordborg *et al.* 2005; Ross-Ibarra *et al.* 2009; Günther and Schmid 2010). Mutations that are annotated as deleterious are particularly likely to occur at lower frequencies than other classes of variants (Marth *et al.* 2011; Kono *et al.* 2016; Liu *et al.* 2017) and may be less likely to be shared among populations (Marth *et al.* 2011).

Tools for the annotation of potentially deleterious variants continue to be developed rapidly (see Grimm *et al.* 2015 for a recent comparison). This includes many tools that attempt to make use of information beyond sequence conservation, including potential effects of variants on protein structure or function (Adzhubei *et al.* 2010) or a diversity of genomic information intended to improve prediction of pathogenicity in humans (Kircher *et al.* 2014). The majority of SNP annotation tools are designed to work on human data and may not be applicable to other organisms (Kono *et al. bioRxiv*). There is the potential for circularity when an annotation tool is trained on the basis of pathogenic variants in humans and then evaluated on the basis of a potentially overlapping set of variants (Grimm *et al.* 2015). Even given these limitations, validation outside of humans is more challenging because of a paucity of known phenotype-changing variants. To address this issue, Kono *et al. (bioRxiv)* report a comparison of seven annotation tools applied to a set of 2,910 phenotype-changing variants in the model plant species *Arabidopsis thaliana.* The authors find that all seven tools more accurately identify phenotype-changing variants likely to be deleterious in *Arabidopsis* (Kono *et al. bioRxiv*) than in humans (Grimm *et al.* 2015; Dong *et al.* 2015). Diversity estimates in *Arabidopsis thaliana* suggest a slightly larger estimated N_e_ in *Arabidopsis* than humans (Cao *et al.* 2011). Given the general principle that for variants with selective coefficients s < 1/(2N_e_), genetic drift will dominate over the action of selection on a variant, making purifying selection less effective.

*No one is perfect, not even the reference*

Bioinformatic approaches can suffer from reference bias (Simons *et al.* 2014; Kono *et al.* 2016; Liu *et al.* 2017). With reference-based read mapping, variants are typically identified as differences from reference, then passed through a series of filters to identify putatively deleterious variants. Most annotation approaches focus on nonsynonymous differences from references. Because a reference genome, particularly when based on an inbred, has no differences from itself, the reference has no nonsynonymous variants to annotate and thus appears free of deleterious variants. This issue can be addressed by identifying all variants, including those where the reference is different from all other samples, and defining the mutation as the change relative to an inferred ancestral state (see Kono *et al.* 2016). A more concerning type of reference bias is that individuals that are genetically more similar to the reference genome will have fewer differences from reference and thus fewer variants that annotate as deleterious. This pattern is observed by Mezmouk and Ross-Ibarra (2014) who find fewer deleterious variants in the stiff-stalk population of maize (to which the reference genome B73 belongs) than in other elite maize populations. Along similar lines, because gene models are derived from the reference genome, more closely related lines with more similar coding portions of genes will appear to have fewer disruptions of coding sequence (Gan *et al.* 2011). Finally, reference bias can contribute to under-calling of deleterious variants, either when divergent haplotypes fail to properly align to the reference or if the reference is included in a phylogenetic comparison. Variants that are detected as a difference from reference may then be compared against the reference in an alignment testing for conservation at a nucleotide. This conflates diversity within a species with the phylogenetic divergence that is being tested in the alignment. In cases where the reference genome contains the novel (or derived) variant compared to the state in the related species, the presence of the reference variant in the alignment will cause the site to appear less constrained. This last problem is easily resolved by leaving the species being tested out of the phylogenetic alignment used to annotate deleterious variants (as in Schubert *et al.* 2014; Henn *et al.* 2016; Kono *et al.* 2016; Marsden *et al.* 2016).

### Effect size

The evidence we present supports the cost of domestication hypothesis, namely that domesticated lineages carry more or higher frequency deleterious variants than their wild relatives (Table 1). However, the distribution of fitness effects of variants is important with the regard to the total load within an individual or population (Henn *et al.* 2015). In other words, it is the cumulative effect of the variants carried by an individual that make up its mutational load, not simply what proportion of those variants is ‘deleterious’. As we describe above, current bioinformatic approaches that rely on phylogenetic conservation may identify a number of false positives (driven by new fitness optima under artificial selection) or false negatives (with specific epistatic, dominance, or environmentally-dependent effects). Functional and quantitative genetics approaches provide means of assessing the phenotypic effect of genetic variants, but there are practical considerations that limit the quantity of evaluations that can be conducted.

One potential issue is that the phenotypic effect of a genetic variant often depends on genomic background, through both dominance (interaction between alleles at the same locus) and epistasis (interaction among alleles at different loci). Evaluating the effect of these variants is consequently a complex task, requiring the creation and evaluation of multiple classes of recombinant genomes. Similarly, the environment is an important consideration in thinking about effect size. As we saw with the yield under drought *qDTY1.1* allele in rice *(Identifying dangerous hitchhikers,* above; Vikram *et al.* 2015), the effect of a genetic variant can depend on developmental or assessment environment.

These kinds of variants might be expected to be involved in local adaptation in wild populations, and so would not show up in a screen based on phylogenetic conservation. Nevertheless, these variants may be important in breeding programs that target general-purpose genotypes with, for example, high mean and low variance yields. Identifying these alleles requires assessment in the appropriate environment(s) and large assessment populations (as the power for detecting genotype-phenotype correlations generally scales with the number of genotypes assessed). Addressing effect size in the appropriate context(s) therefore involves challenges of scale. Fortunately, new types of mapping populations (e.g. MAGIC) and high-throughput phenotyping platforms that can enable this work are increasingly available across domesticated systems. Given these tools and the resultant data, we should soon be able to parameterize genomic selection and similar models with putatively deleterious variants and test their cumulative effects.

### Heterosis

In systems with hybrid production, complementation of deleterious variants between heterotic breeding pools may contribute substantially to heterosis (e.g. in maize; Hufford *et al.* 2012; Mezmouk and Ross-Ibarra 2014; Yang *et al.* bioRxiv). If this is broadly true, hybrid production may be an interesting solution to the cost of domestication, as long as deleterious variants are private to heterotic groups rather than fixed in domesticated species. Theory suggests that the deleterious variants that contribute to heterosis between populations with low levels of gene flow are likely to be of intermediate effect, and may not play a large role in inbreeding depression (Whitlock, Ingvarsson, and Hatfield 2000). Since fitness in these contexts is largely evaluated in hybrid individuals rather than inbred parents, parental populations are likely to retain a higher proportion of slightly and moderately deleterious variants than even selfing populations (Figure 2A). As long as hybrid crosses are the primary mode of seed production, these alleles may not be of high importance to breeders even though they contribute to deleterious variation more broadly. Evaluating effect size in this context requires assessing both parental and hybrid populations, and is therefore that much more difficult. However, assuming that complementation of deleterious variants is a substantial component of heterosis, quantifying these variants should improve the ability of breeding programs to predict trait values in hybrid crosses. Further, the question of whether deleterious variation limits genetic gains in hybrid production systems remains open.

## CONCLUSIONS

Research on the putative costs of domestication is still relatively new, and there remain many open questions. However, our review of the literature suggests that deleterious variants are generally more numerous or frequent in domesticated species compared to their wild relatives. This pattern is likely driven by a number of processes that collectively act to reduce the efficacy of selection relative to drift in domesticated populations, resulting in increased frequency of deleterious variants linked to selected loci and greater accumulation of deleterious variants genome-wide. We encourage further research across domesticated species on these processes, and recommend that researchers: (1) Sample domesticated and wild lineages sufficiently to assess diversity within as well as between these groups and (2) present deleterious variant data per genome and as proportional as well as absolute values. We also strongly encourage researchers and breeders to think about deleterious variation in context, both genomic and environmental. Finally, we think collecting empirical data on the effect sizes and conditional dependence of putatively deleterious variants is increasingly feasible, and would contribute greatly to this field.

## ACKNOWLEDGEMENTS

We thank Greg Baute, Thomas Kono, Kathryn Turner, Jeffrey Ross-Ibarra, and two anonymous and thoughtful reviewers for comments on an earlier version of this manuscript, and Brandon Gaut, Loren Rieseberg, and Greg Owens for helpful discussion.

## Supplementary Table 1

Quantitative version of Table 1: evidence for a cost of domestication across domesticated plants and animals, with *Arabidopsis thaliana* and *Homo sapiens* included for comparison. Plant 1C genome sizes from the RBG Kew database (http://data.kew.org/cvalues/), except tomato (Michaelson *et al.* 1991) and *Cynara* spp. (Giorgi *et al.* 2016). Plant chromosome counts from the Chromosome Counts Database (http://ccdb.tau.ac.il/). Animal 1C genome sizes and chromosome counts from the Animal Genome Size Database (http://www.genomesize.com/, mean value when multiple records were available). Gene numbers are high-confidence (if available) estimates from the vertebrate and plant Ensembl databases (http://uswest.ensembl.org/; http://plants.ensembl.org/), except common bean (Schmutz et al. 2013), sunflower (Compositate Genome Project, unpublished), and *Cynara* spp. (Scaglione et al. 2016). LD N50 is the approximate distance over which LD decays to half of maximum value. Loss of genetic diversity is calculated as 1 – (ratio of p_domesticated_ to p_wild_), or q if p not available.

